# Anti-angiogenic activities of the leaves of *Tapinanthus bangwensis* (Engl. and K. Krause) Danser Loranthaceae growing on three host trees in south-western Nigeria

**DOI:** 10.1101/759209

**Authors:** Oluwatofunmilayo Arike Diyaolu, Alfred Francis Attah, AS Adeagbo, Y Fukushi, Jones Olanrewaju Moody

## Abstract

The control of angiogenic process is of immense importance to the overall health of mankind to avert out-of-control complications such as tumours and cancers. *Tapinanthus bangwensis* (TBG) is a semi-parasitic plant with multipurpose uses in Nigerian ethno-medicine among which is its anti-infective properties as well as the anti-tumor and restoration of damaged tissues. Despite the wide and varied uses of TBG, the anti-angiogenic investigation of the plant on different host trees is lacking while the documentation and scientific validation of indigenous knowledge on plants’ use is urgent. The present work focuses on the anti-angiogenic activities of the crude extracts and partitioned fractions of the leaves of *Tapinanthus bangwensis* (TBG) using the Chick chorio allantoic membrane (CAM) model. The plant was collected from three different host trees namely *Albizia lebbeck* (AL), *Stereospermum kunthianum* (SK) *and Tabebuia rosea* (TR). The methanolic crude extracts and solvent-partitioned fractions of TBG-SK samples were assessed for their anti-angiogenic activities using the chick chorio allantoic membrane (CAM) *in ovo* and *in vitro* assay methods respectively. Purification and isolation of major compound(s) in the chloroform fraction of TBG extract obtained from *Stereospermum kunthianum* host tree (TBG-SK) was carried out using chromatographic and spectroscopic methods (NMR, MS).

The crude methanolic extract of TBG-SK was most potent (100% activity) in the CAM *in ovo* assay while the chloroform fraction produced a significantly (p<0.05) highest average reduction in blood vessels with resultant formation of large avascular zones on CAM following an *in vitro* CAM assay. The anti-angiogenic chloroform fraction revealed the presence of a UV-active triterpenoid moiety. Findings from this work, has provided some justification for the folkloric use of TBG and thus forms a potential basis for drug discovery for wide-ranging disease states.

## Introduction

Angiogenesis is the proliferation new blood vessels from an existing one. It is a well controlled process that rarely occurs under normal conditions, except for instances of wound healing, embryonic development and formation of the corpus luteum [1]. The success of angiogenesis depends on the sensitive equilibrium between growth stimulating and inhibiting factors [2]. In many serious disease states the body loses control over angiogenesis. Angiogenesis-dependent diseases result when new blood vessels either grow excessively or insufficiently. For instance excessive angiogenesis occurs in diseases such as cancer, diabetic blindness, age-related macular degeneration, and rheumatoid arthritis while insufficient angiogenesis can be seen in diseases such as coronary artery disease, stroke, and chronic wounds [3].

Angiogenesis has become a promising target for experimental therapies in cancer and many other life-threatening diseases. Wide varieties of therapies directed at interfering with this process are in development. A number of clinical drugs such as Bevacizumab (Avastin^®^) have been developed to combat excessive angiogenesis in cancers [4]. However, major hurdles for clinical implementation include limited efficacy, rapid development of resistance to the anti-angiogenic modalities and in some cases severe toxicity, arising from growth factors [5]. Hence the need for identification of new drugs particularly plant-derived agents that are non-toxic and effectively act at various steps of angiogenic cascade. Nature-derived drug discovery effort targeting angiogenesis could be of great clinical significance. This is well supported by the reported gradual drift from the use of synthetic drugs to plant-based supplements and remedies [6]. Interestingly, reports from the World Health Organisation say at least 80% of populations in developing world depend on plants for their primary healthcare needs [7]. Many plants from the Nigerian ethnomedicine have been sources of diverse therapeutic agents [8]. However, only a few of them have been scientifically evaluated for their angiogenesis modulatory activity.

*Tapinanthus bangwensis* (TBG) commonly known as the African mistletoe is a semi-parasitic, semi-ligneous plant, 1-2 cm long, arranged in tufts, with drooping branches. It consists basically of globular haustoria which fit into a socket of hypertrophied host tissues to form an irregular fused ball and socket structure. It grows on different host trees [9]. The African mistletoe is an important plant in Nigerian ethnomedicine. Extracts from the leaves are used by traditional medical practitioners to treat various ailments. The mistletoe plant has been described as “an all purpose herb” because of its rich folkloric uses in the management of several disease conditions [10]. However, there is no documented scientific data on the angiomodulatory effect of the plant. This study was therefore designed to assess the effect of extracts of TBG growing on three host plants, on angiogenesis. Previous phytochemical and bioactivity studies on *the plant* have revealed the presence of a variety of secondary metabolites [11]. According to a recent review new triterpenoids have been isolated from the seeds [12] and a variety of bioactivities such as anticancer, anti-inflammatory, anti-oxidant, antimicrobial and anti-diabetic activities have been reported [13, 14]. In this study, we have screened the ability of TBG to inhibit or promote angiogenesis using the chick chorio alantoic membrane *in ovo* and *in vitro* assays.

## 2. Materials and methods

## 2.1 General

The study was carried out in line with the NIH standard of care and good research practice. This experiment did not involve rodents or other experimental animals which requires ethical approval from the University of Ibadan animal welfare and ethics committee. However, avian fertilized eggs were used and at the time of the study, no ethical approval or waiver was required except the University general research policy which we carefully adopted by ensuring that all experiments involving life at any stage were handled humanely in line with the internationally accepted standards for conducting laboratory experiments.

The number of blood vessels was counted using a stereomicroscope (Model SM-5 stereoscopic microscope).

The proton NMR experiment was performed on a 270 MHz spectrometer (JEOL GX-270 MHz Fourier Transform Nuclear Magnetic Resonance 6.3 Tesla Spectrometer) and referenced to residual solvent peaks (CDCl_3_: *δ*_H_ 4.17, *δ*_H_ 5.20, *δ*_H_ 2.36, *δ*_H_ 1.99, *δ*_H_ 0.85, *δ*_H_ 0.79, *δ*_H_ 0.87, *δ*_H_ 0.96, *δ*_H_ 1.06, *δ*_H_ 1.04). Mass spectra data was acquired using an instrument with configuration: FD, MS-T100GCV; Ionization Mode: FD^+^ (Field desorption).

Column chromatography (CC) was performed on silica gel (230–400 mesh) using a column of length 60 cm and internal diameter of 3.5 cm. All solvents used for CC were of analytical grade. Pre-coated silica gel GF_254_ plates were used for thin-layer chromatography and preparative thin-layer chromatography (TLC) analyses.

## 2.2 Plant collection, extraction and preliminary chemical screening

The leaves of TBG were collected from three different host trees namely: *Albizia lebbeck, Stereospermum kunthianum* and *Tabebuia rosea* in April, 2013 from the Botanical Garden, University of Ibadan. The plant was authenticated at the Forest Herbarium Ibadan, where voucher specimen, FHI 109822 was deposited. Preliminary phytochemical screening of powdered leaves was carried out using standard protocols reported by Trease and Evans [15] to detect the presence of alkaloids, tannins, flavonoids, cardiac glycosides, saponins, terpenoids, anthraquinones and steroids. The air-dried leaf powder of TBG collected from the three host trees code named TBG-AL, TBG-SK and TBG-SR were subjected to solvent extraction using methanol under the room temperature (around 25 °C). Extract obtained was kept referigerated at 4 degrees for a selective preliminary biological screening. The air-dried powder of the leaves of TBG growing on *Stereospermum kunthianum* (TBG-SK) (2 kg), being the most biologically active, was extracted with absolute methanol at room temperature. Filterate obtained was concentrated in-vacuo and the yield of crude extract determined. Dried crude extract (30 g) was further subjected to exhaustive liquid-liquid partitioning using chloroform, ethyl acetate and aqueous methanol. The chloroform fraction (9 g) was submitted to column chromatography (CC) over silica gel (230–400 μm) applying a 10% gradient of increasing polarity with hexane, ethyl acetate and methanol. Around 50 mL each of 122 fractionated eluates were collected and pooled together by similarity of TLC spots leading to a total of 21 fractions (Cb-1 to Cb-21). The fraction Cb-2 (3 mg) was subjected to preparative thin layer chromatography (Prep-TLC) using pre-coated silica gel GF_254_ plates with CHCl_3_: EtOAc (9:1).

### 2.3 CAM in ovo assay

Day old fertilized Isa brown chicken (*Gallus domesticus*) eggs “Fig 1A” were procured from the Ona ara local government hatchery. They were washed with tap water to remove any dirt. The eggs were then incubated at 37 °C and 70 % relative humidity “Fig 1B”. The eggs were rotated manually twice a day during incubation to prevent embryo from adhering to the shell membrane. After 72 h, the eggs were examined by candling (illuminating the egg shell with a 60 W bulb) [6] “Fig 1C”. This confirms the viability of the embryo. The eggs were then wiped with 70% ethanol, and kept in an upright position with the blunt end upwards to allow embryo alignment with the air sac [16]. A small aperture (4 mm diameter) was made at the narrow end of the egg using pointed scissors and 2 mL of albumen was removed “Fig 1D”. The hole was sealed with sterile cotton ball and sealing tape. A window of 2 cm × 2 cm was then cut on the blunt end and the pieces of egg shell were carefully removed using sterile forceps. The inner shell membrane was then cut to expose the embryo and the CAM vasculature for the application of test substances “Fig 1E”. Dried sterile Whatman filter paper No. 2 disc loaded with the crude extracts and positive control (Itraconazole) at different strengths (0.5, 1.0 and 1.5 mg/mL) was then placed on the CAM vasculature. This was done in triplicate and untreated CAM’s served as the master control. The number of eggs per group was 5. The window was closed using sealing tape and the eggs were returned to the incubator for another 48 h at 37°C with a relative humidity of 70 %. After incubating for 48 h, the egg shells were cut open to expose the complete expanse of CAM vessels and to examine the extent of inhibition of blood vessel growth. Images of the CAMs were captured using a Huawei G700 camera for comparative studies. The most active extract was then selected for further studies.

**Fig. 1A:**
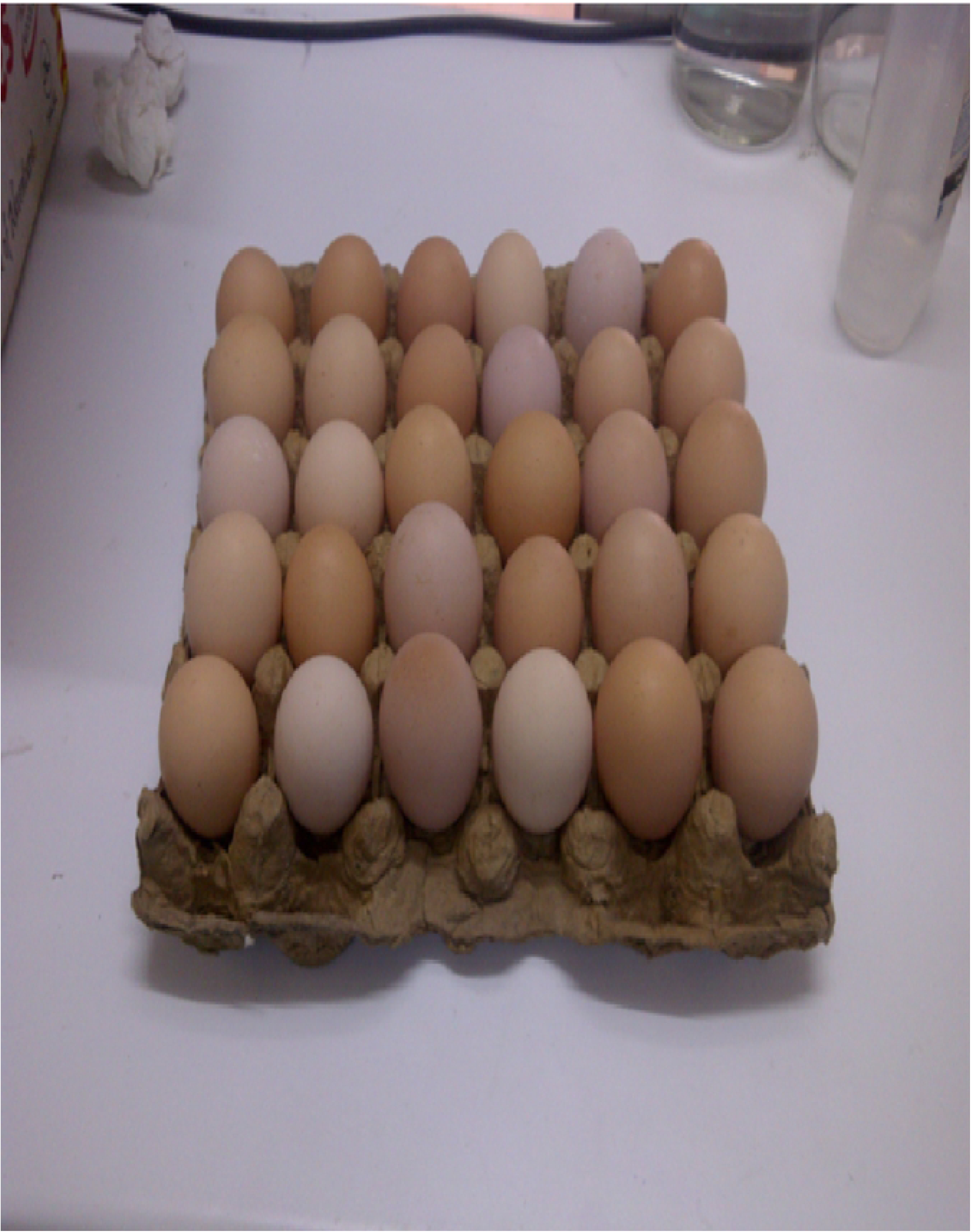
Freshly fertilized eggs(*Gallus domesticus*)

**Fig. 1B:**
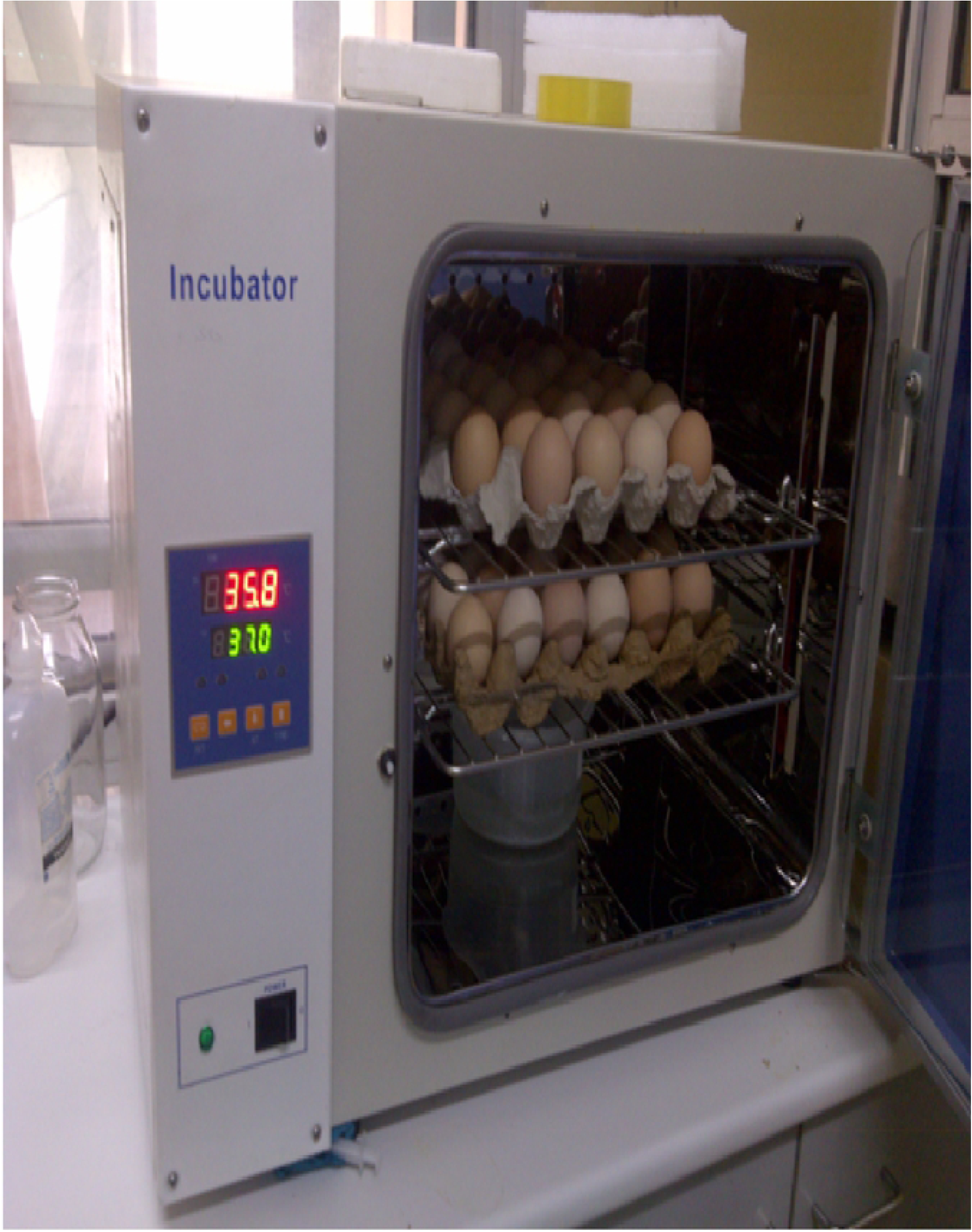
Eggs were incubated at 37 °C and 70 % humidity

**Fig. 1C:**
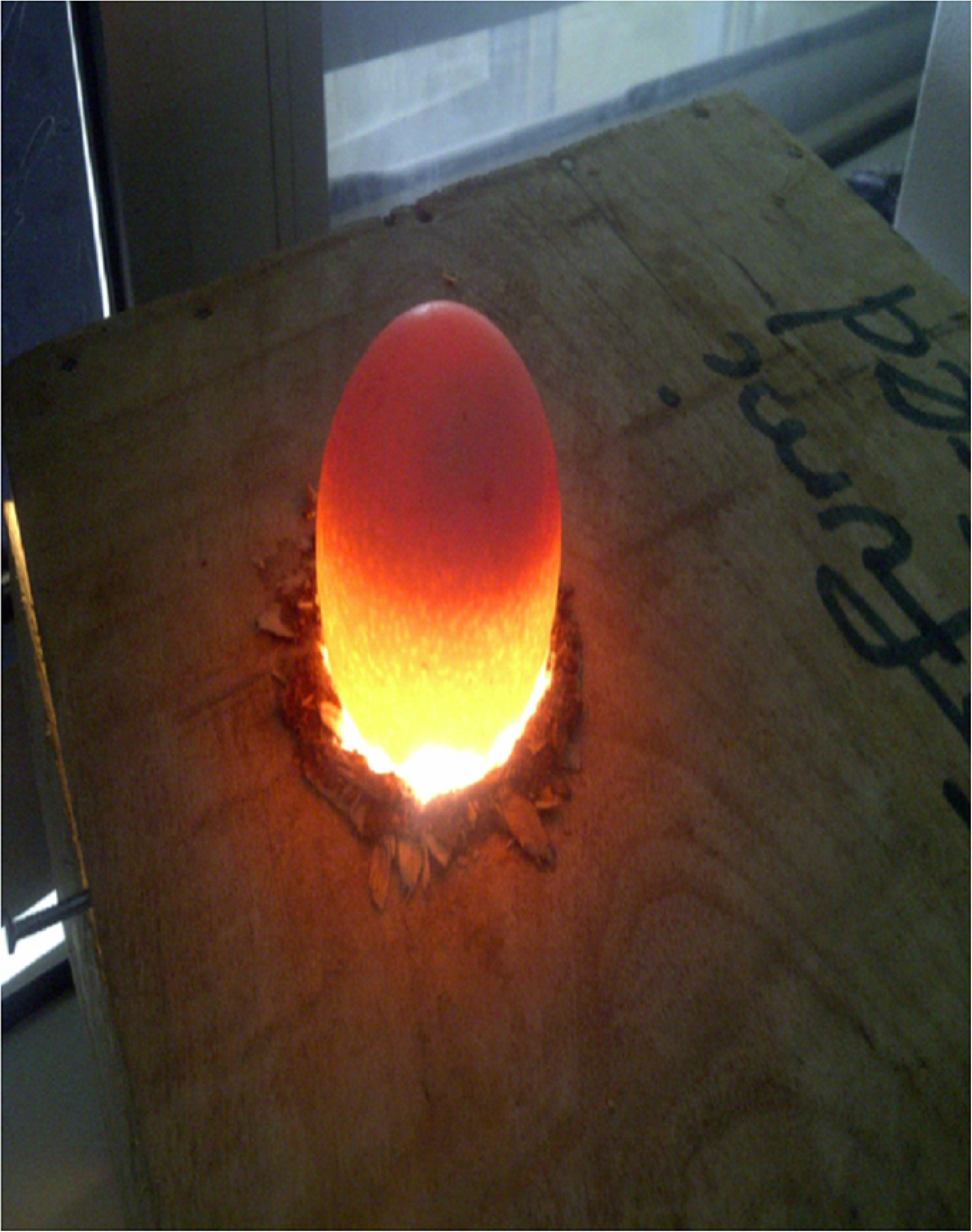
Eggs were candled to ascertain viability

**Fig. 1D:**
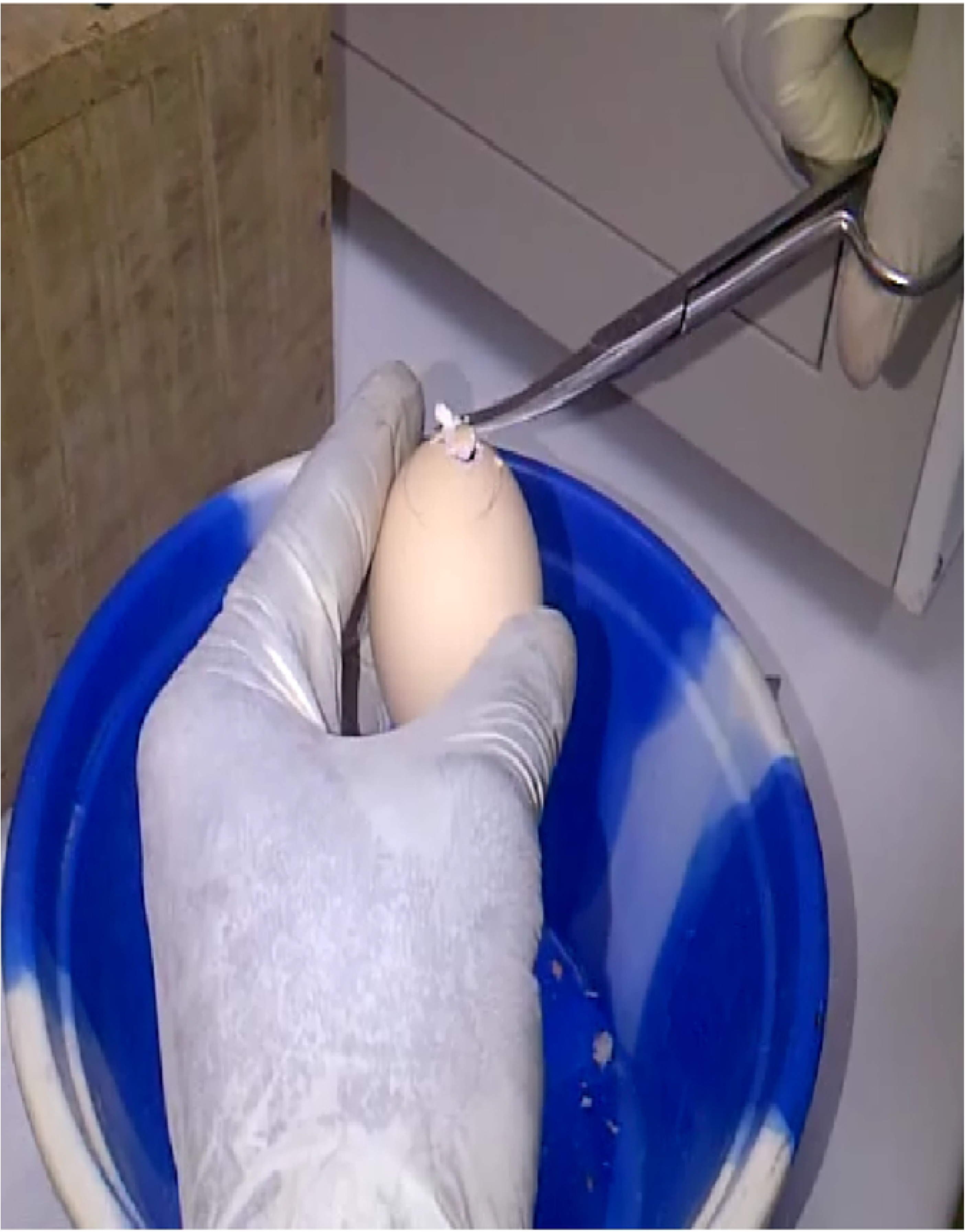
Removal of thin albumin

**Fig. 1E:**
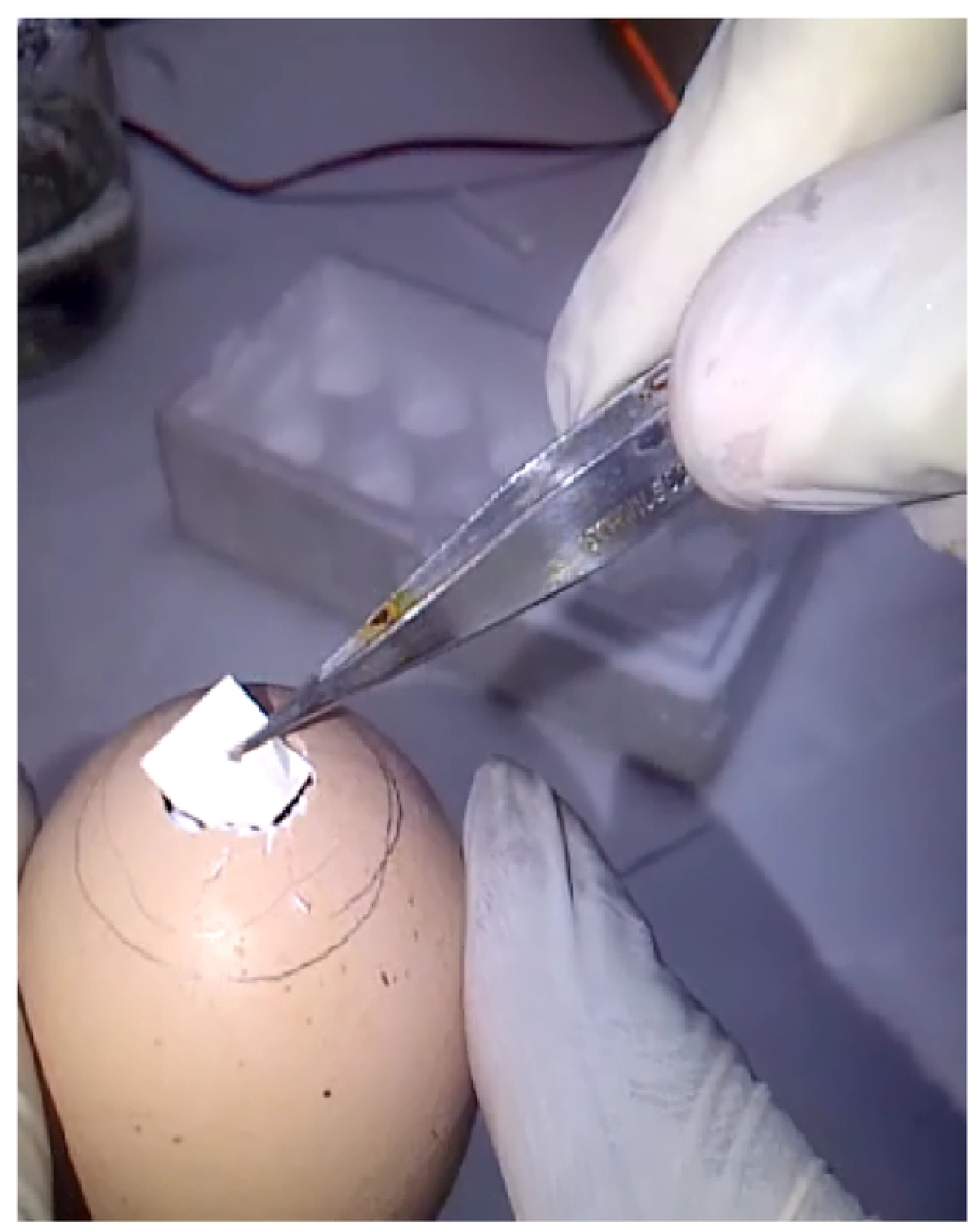
Samples to be tested was applied on the vessels

**Fig. 1F:**
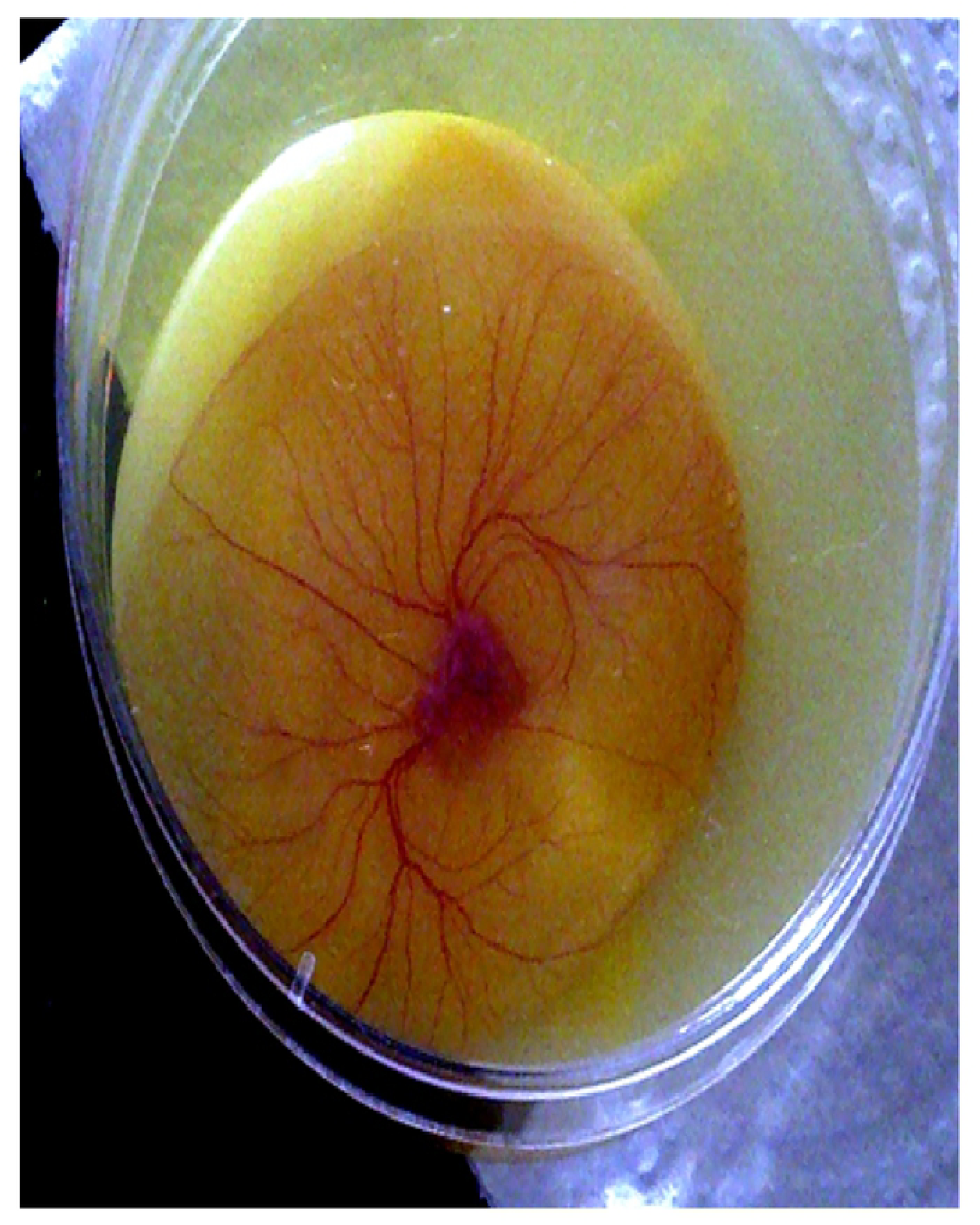
In vitro assay provides enlarged observation field for quantitation

### 2.4 CAM *in vitro* anti-angiogenic assay

CAM assay was performed as previously described with some modifications. Freshly fertilized Isa brown chicken (*G. domesticus*) eggs procured from Ona ara local government hatchery were washed with tap water to remove dirt. Eggs were then incubated for 3 days in an incubator at 37 °C and 70 % relative humidity. The eggs were rotated manually twice a day during incubation to prevent embryo adherence to the shell membrane. On the 3rd day of incubation, eggs were candled to confirm the development stage and viability. The eggs were divided into five test groups: The positive control (Itraconazole 0.25mg), master control (untreated CAM’s), chloroform fraction (A), ethylacetate (B) and aqueous methanol (C) test group (fractions obtained from the liquid-liquid partitioning). Each group comprised ten eggs and the experiment was carried out in triplicate. The eggshell was cracked and gently opened into a sterile petri dish “Fig 1F”. It was made sure that the yolk sac membrane remained intact and that the embryo was viable. The number of blood vessels in each Petri dish was counted using a stereomicroscope. Sterile whatman filter paper No. 2 disc loaded with the different test fractions and allowed to dry under sterile conditions at different concentrations (0.25, 0.50 and 0.75 mg/mL) was placed in areas between vessels and incubated for another 24 h. The number of blood vessels remaining after the incubation period was counted again “Fig 2”. Results of the samples obtained were photographed using a Huawei G700 camera for comparative studies and quantitation.

**Fig 2.**
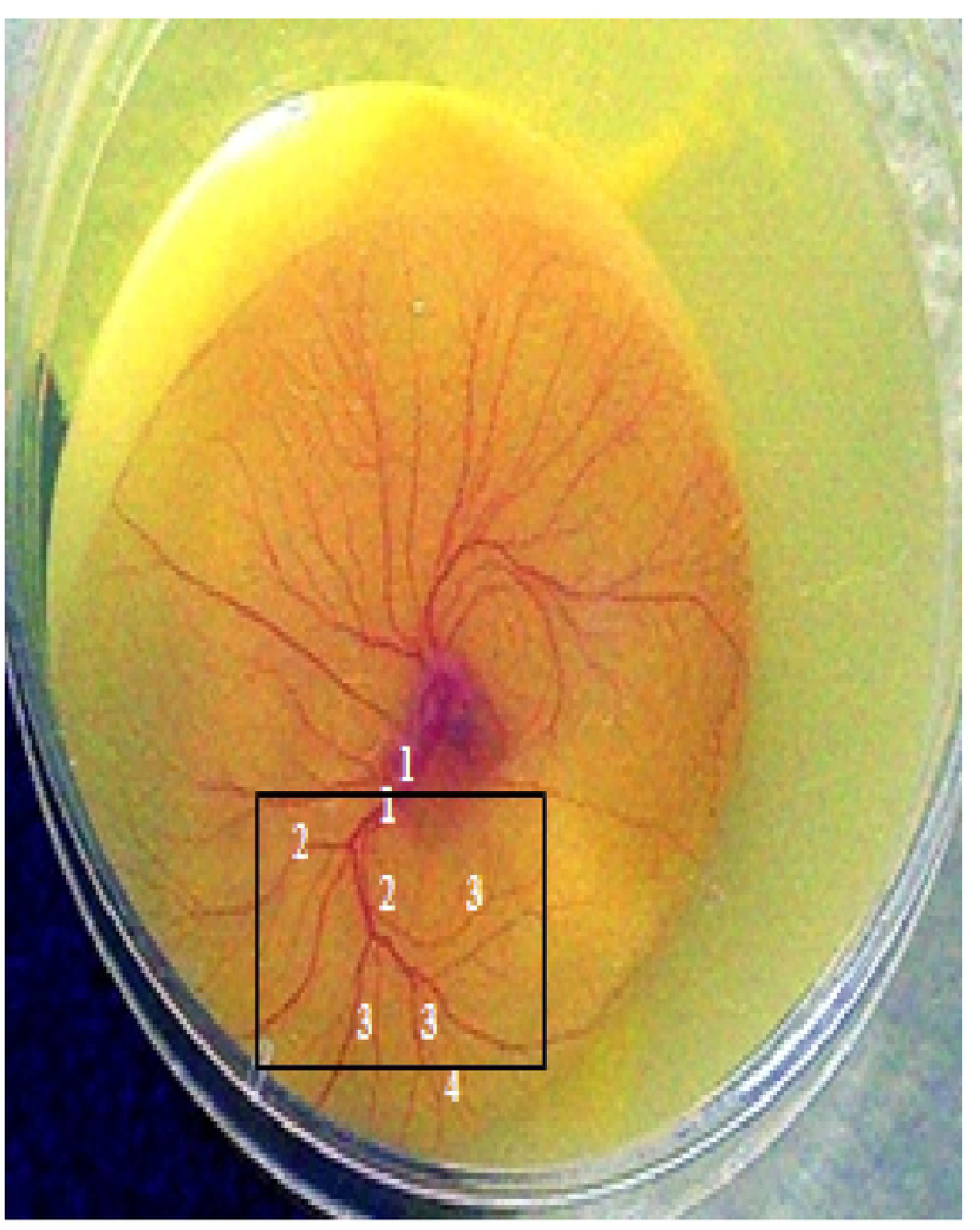
Counting blood vessels in *in vitro* CAM. 1: primary blood vessels; 2: secondary blood vessels; 3: tertiary blood vessels; 4: quaternary blood vessels.

### 2.5 Thin layer chromatography

Thin layer chromatography screening was done on the chloroform fraction of TBG-SK. Samples were applied to silica gel 60 F_254_ chromatographic plates at the same delivery speed, with a distance of 10 mm between them, and the 10 mm distance from all edges of the plate. Different eluent systems were used: chloroform: methanol = 9:1 (v/v)(mobile phase I), hexane: ethylacetate = 8:2 (v/v)(mobile phase II) and toluene: acetone: ethylacetate = 8:1:1(v/v/v)(mobile phase III). The plates were developed over a path of 5 cm and air dried. All plates were examined under daylight and with UV light (λ= 254nm and 365 nm) (Camag Reproster lamp). The plates were then sprayed with: anisaldehyde in sulphuric acid, vanillin in sulphuric acid and ferric chloride.

### 2.6 Counting of blood vessels

A black square of size 150 × 150 pixels was placed on areas where the effect of test samples was observed. The number of primary, secondary, tertiary, and quaternary blood vessels was counted using the stereomicroscope “Fig 2”.

### 2.6 Statistical analysis

Data were presented as mean ± standard error of mean (SEM). Results were calculated by using the statistical software (SPSS, version 13.0, SPSS Inc., Chicago, USA).

## 3. Results and discussion

The chick chorio allantoic membrane (CAM) is an extra embryonic tissue consisting of two fused epidermal layers (chorion ectoderm and allantoic endoderm) that enclose a mesodermal double layer in which a dense vascular network is formed [17]. In this assay, withdrawing 2 mL of albumen allows for easy opening of the egg shell without causing any physical stress to the embryo. There was no contamination during the experiment.

TBG-AL had the least anti-angiogenic activity showing 100%. 93.3% and 80% survival of live embryos at the respective concentrations. TBG-TR extract showed a moderate increase in activity with a corresponding increase in dose administered, gradually decreasing the survival percentage from 100% to 66.7% “Fig 3”.

**Fig 3.**
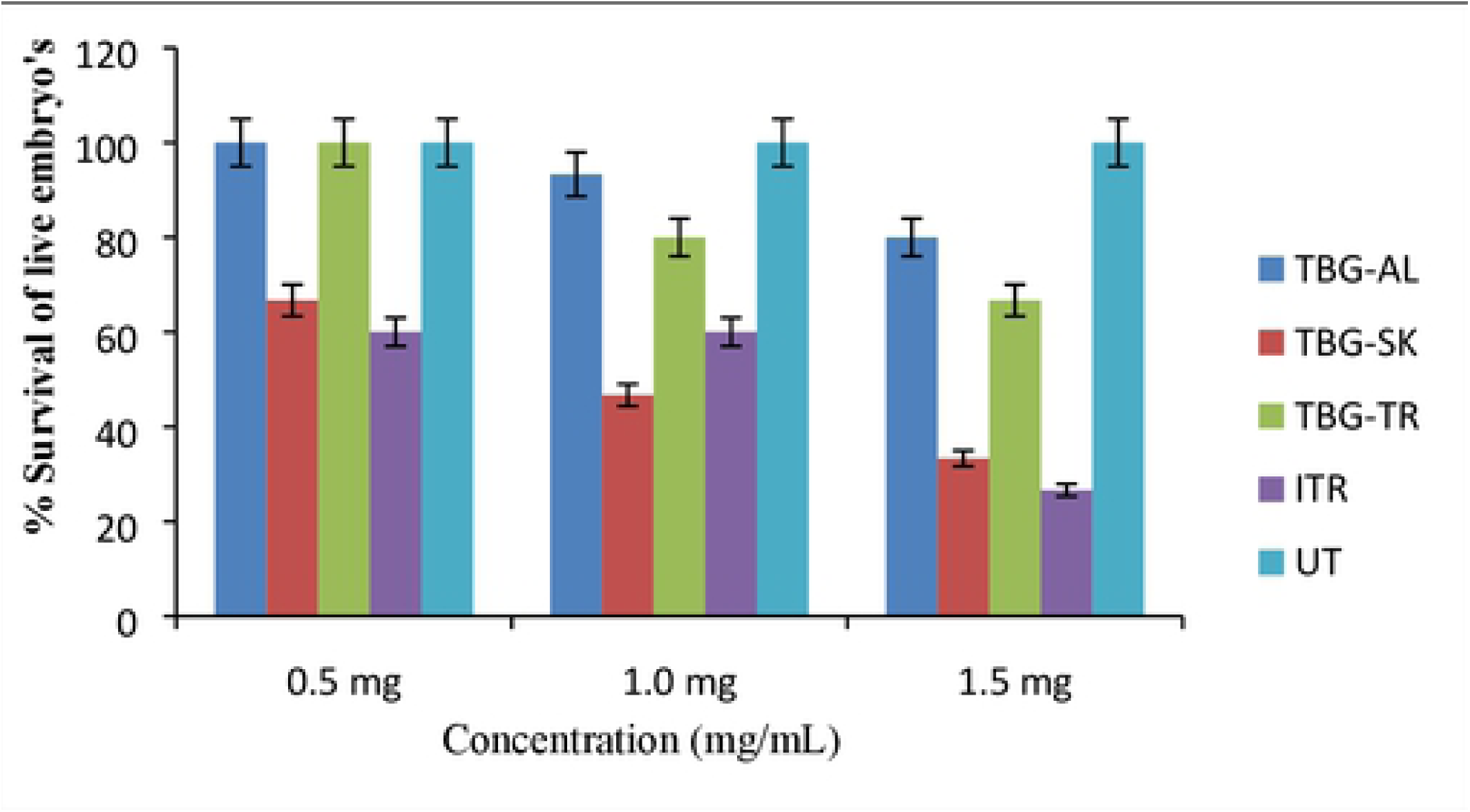
Percentage survival of living embryos following treatment with TBG extracts and Itraconazole. TBG-AL: *Tapinanthus bangwensis* growing on host *Albizia lebbeck* TBG-SK: *Tapinanthus bangwensis* growing on host *Stereospermum kunthianum* TBG-TR: *Tapinanthus bangwensis* growing on host *Tabebuia rosea*; ITR: Itraconazole UT: Untreated CAM’s

TBG-SK had the highest anti-angiogenic activity, also gradually increasing with an increase in dose. It was thus selected for further studies. The percentage of survival for Untreated CAM’s was constant throughout the experiment, confirming that there was no contamination. The positive control used was itraconazole dissolved in dimethyl sulphoxide (DMSO). Itraconazole is an anti-fungal drug that has been shown to have potent anti-angiogenic activity and is currently used as an adjunct in the management of lung, skin and prostate cancer [18]. The choice of this drug as a standard in this experiment is therefore based on its efficacy, ease of availability and cost. Itraconazole showed the same inhibition rate at 0.5 and 1.0 mg. One will expect an increase in mortality with an increase in concentration. But this lack of increase in activity may be due to the variation in the sizes of blood vessels present in the embryo. The CAM *in vitro* model is better than the pilot *in ovo* model in that; It allows room for quantitation. The number of blood vessels before and after inoculating the test samples into the live embryo can be counted with the aid of a stereomicroscope “Fig 2”.

It allows one to observe the live embryo with a beating heart and blood vessels clearly with the naked eye before the experiment and after the experiment, the avascular zone “Fig 1F”.

It allows for snapshots of the blood vessels. It has a shorter period of incubation after inoculation (24 h), thereby allowing one to observe the actual reduction in blood vessels instead of mortality. However, one major drawback of this model is the higher risk of infection due to the fact that the entire embryo is exposed in a Petri-dish, unlike the *in ovo* model where the embryo remains intact within the egg shell throughout, until the end of the experiment when it is cracked open.

Following the CAM *in vitro* assay, all the TBG-SK test samples (chloroform, ethyl acetate and aqueous methanol fractions) were active showing a reduction in the number of blood vessels “Fig 4”.

**Fig 4.**
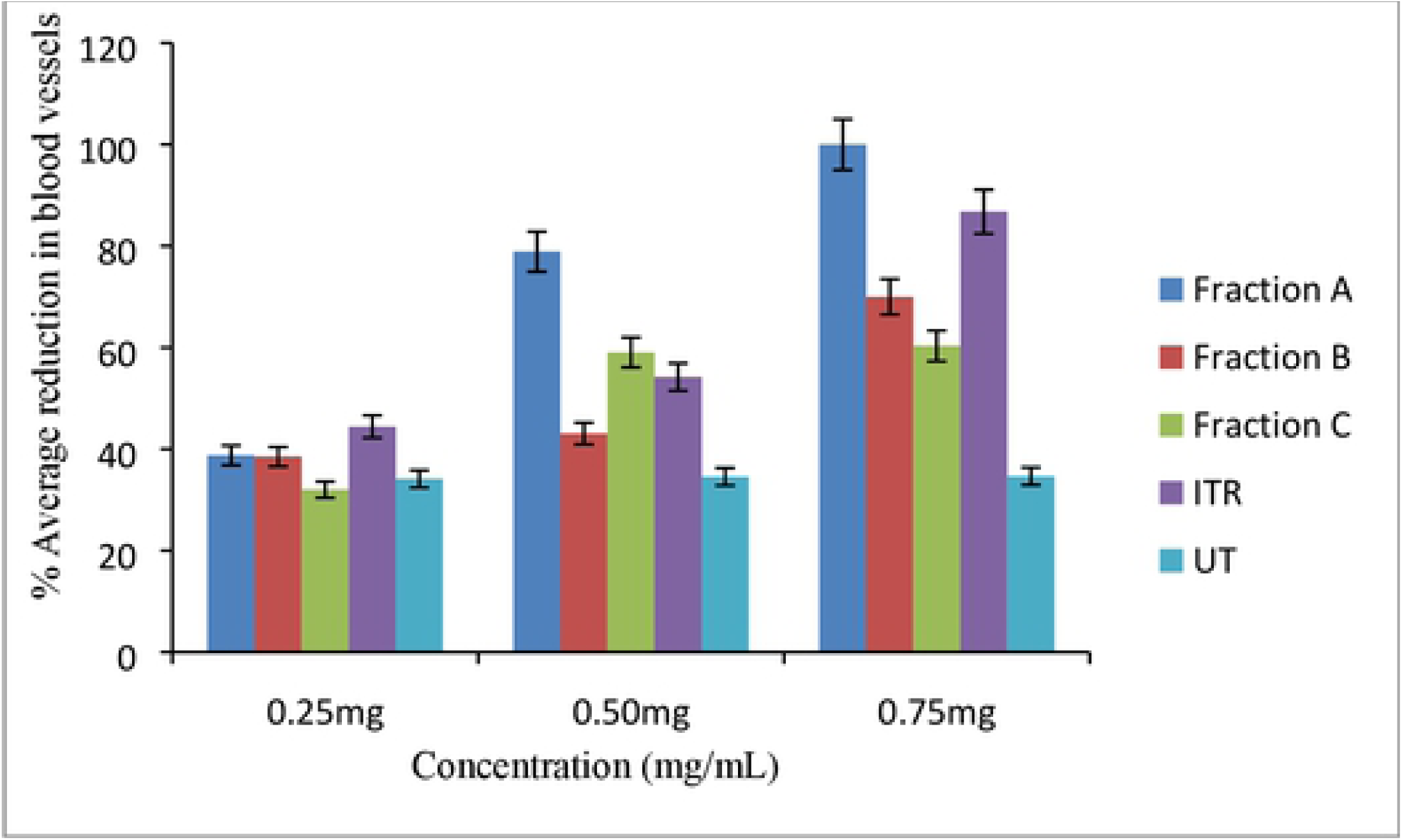
Anti-angiogenic activity of solvent partitioned fractions of TBG-SK. Fraction A: Chloroform fraction; Fraction B: Ethylacetate fraction; Fraction C: Aqueous methanol fraction

The positive control used was 0.25mg/mL Itraconazole dissolved in DMSO. Out of all three fractions screened, fraction A had the highest anti-angiogenic activity, showing a marked reduction in the average number of blood vessels thereby resulting in the formation of large avascular zones “Fig 4”. This activity increased with a corresponding increase in concentration. Fraction B showed moderate activity while fraction C had the least activity. The positive control (Itraconazole 0.25mg/mL) was also very active showing a drastic reduction in neovascularisation in the CAM assay. The Untreated embryo remained exactly the same. i.e. there was no change in the number of blood vessels before and after the experiment.

The preliminary phytochemical screening “Table 1” revealed the extent of host tree influence on the chemical constituents found in the respective mistletoe. These variations indicate that the same species occurring on different hosts in the same locality have differences in their metabolites. The leaves of TBG-SK tested negative to the presence of alkaloids while that of TBG-AL and TBG-TR showed that they contain alkaloids. Also, TBG-TR tested negative to saponins while TBG-SK and TBG-AL tested positive. The differences noted in the chemical constituents of this plant growing on different hosts might justify why the host is as important as the parasite and why the ethnomedicinal use of this parasitic plant in the treatment of an ailment is usually dependent on a particular or specific host.

**Table 1.** Preliminary phytochemical screening

| Sample<br>Code | Part<br>used | Secondary metabolites detected |  |  |  |  |  |  |  |
| --- | --- | --- | --- | --- | --- | --- | --- | --- | --- |
|  |  | Alkaloids | Anthraquinones | Cardiac | glycosides<br>Flavonoids | Saponins | Steroids | Tanins | Terpenoids |
| TBG-AL | Leaf | + | - | + | + | + | + | + | + |
| TBG-SK | Leaf | - | - | + | + | + | + | + | + |
| TBG-TR | Leaf | + | - | + | + | - | + | + | + |

- : absent; + : present; TBG-AL: *Tapinanthus bangwensis* growing on host *Albizia lebbeck*; TBG-SK: *Tapinanthus bangwensis* growing on host *Stereospermum kunthianum*; TBG-TR: *Tapinanthus bangwensis* growing on host *Tabebuia rosea*

Fraction A from TBG-SK being the most active was further purified using column chromatography and TLC. The purification of fraction A yielded sub-fraction CB which was subjected to preparative thin layer chromatography. The developed TLC plates for fraction A were sprayed with different reagents for further analysis of the component compounds. Spraying with anisaldehyde in sulphuric acid produced three main colours: purple, red and orange spots which were all colours that suggest the presence of terpenes or steroidal compounds “Fig 5A-C”. This was eventually confirmed by the results obtained from the proton NMR and mass spectrometric analysis carried out on the isolated compound which revealed the presence of triterpenoid compounds. Ferric chloride spray produced mostly brownish-green coloured spots indicating the presence of tannins [19]. Compound A was obtained as a white amorphous powder as one of the eluates from the column chromatography experiment of fraction A. The FDMS analysis showed a molecular ion peak at *m/z* 426.3877 (calculated 426.3862), corresponding to a molecular formula of C_30_H_50_O with six degrees of unsaturation. The ^1^HNMR spectrum displayed signals for tertiary methyl groups at δ_H_ 0.85(3H, s, CH_3_), 0.79 (3H, s, CH_3_), 0.87 (3H, s, CH_3_), 0.96 (3H, s, CH_3_) 1.06 (3H, s, CH_3_), 1.04 (3H, s, CH_3_) which are characteristic of triterpenoid compounds at the upfield region of the spectral. Two multiplets at δ_ppm_ 2.36 (2H, t, CH_2_) and 1.99 (2H, m, CH_2_), both integrated for two protons each, were ascribed to methylene protons. A one proton singlet at δ_H_ 4.17 (1H, s, OH-3), a more downfield region of the spectra due to the presence on an electronegative atom was ascribed to an hydroxyl group most likely at position 3 of the triterpenoid nucleus. A one proton multiplet occuring at δ_ppm_ 5.20 (1H, m, olefinic – H) in the downfield region which showed a deshielded proton is ascribable to an olefinic proton.

**Fig. 5A:**
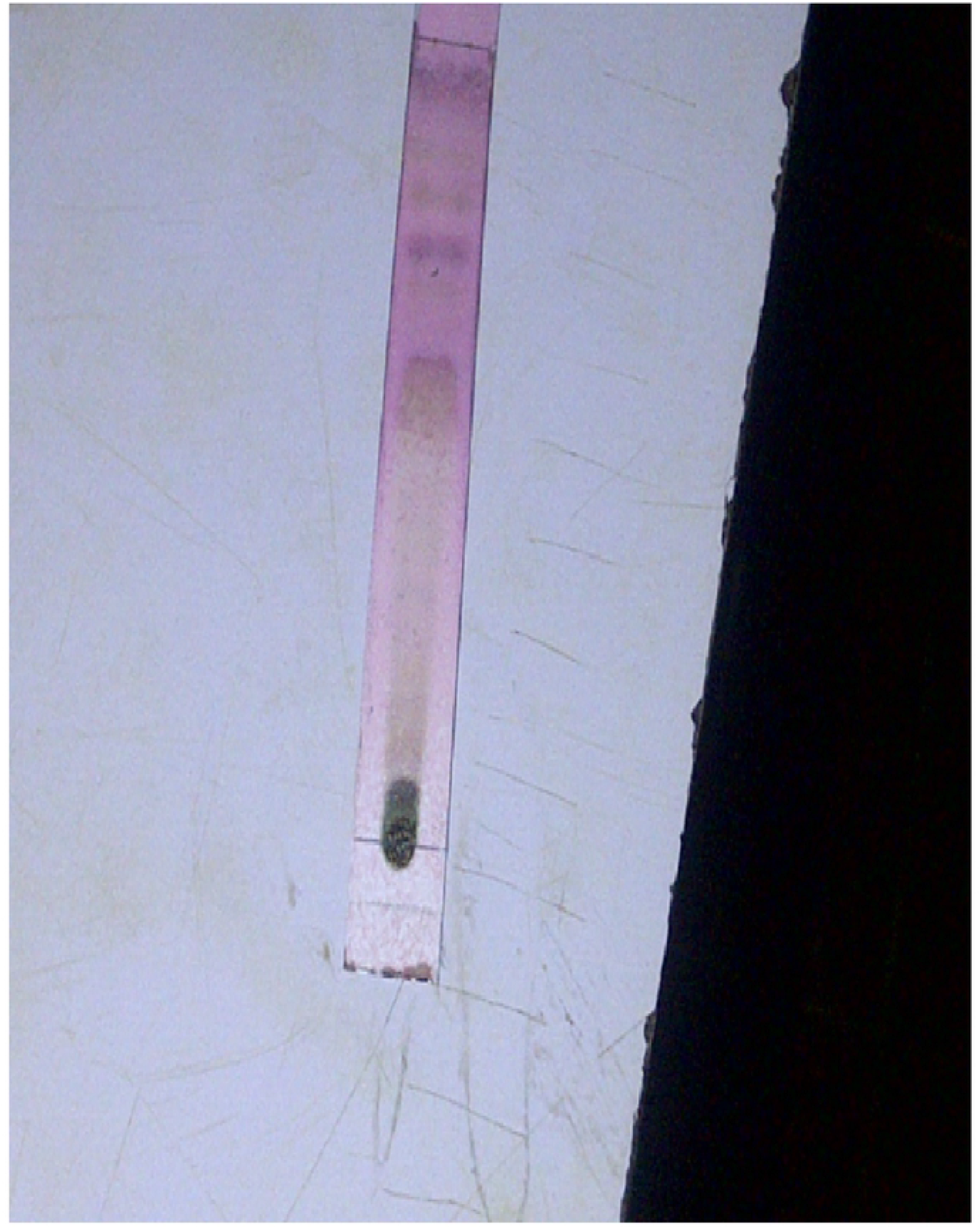
Thin layer chromatographic analysis of chloroform fraction (TBG-SK) sprayed with vanillin in sulphuric acid

**Fig. 5B:**
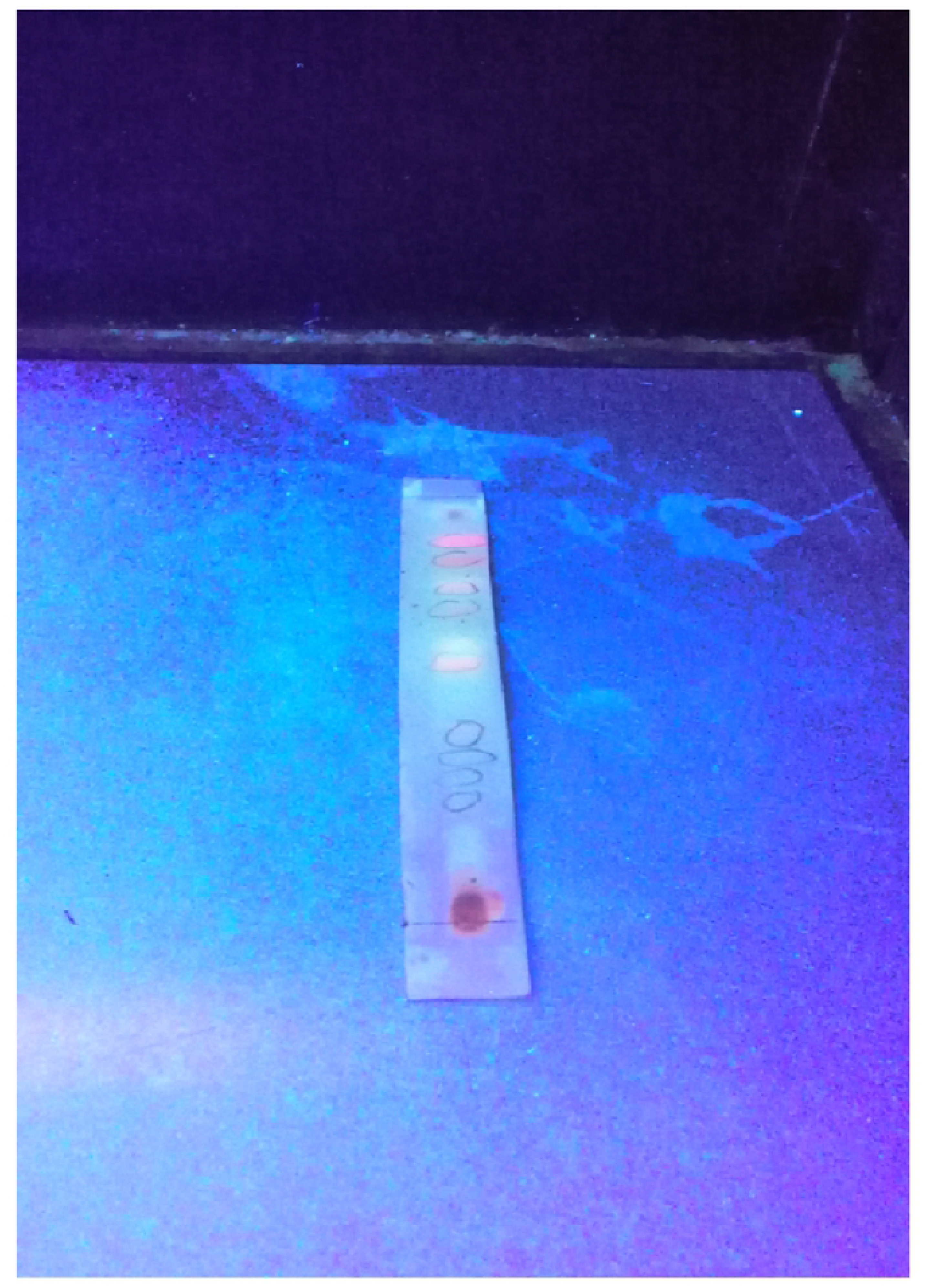
Thin layer chromatographic analysis of chloroform fraction (TBG-SK) sprayed with anisaldehydein sulphuric acid

**Fig. 5C:**
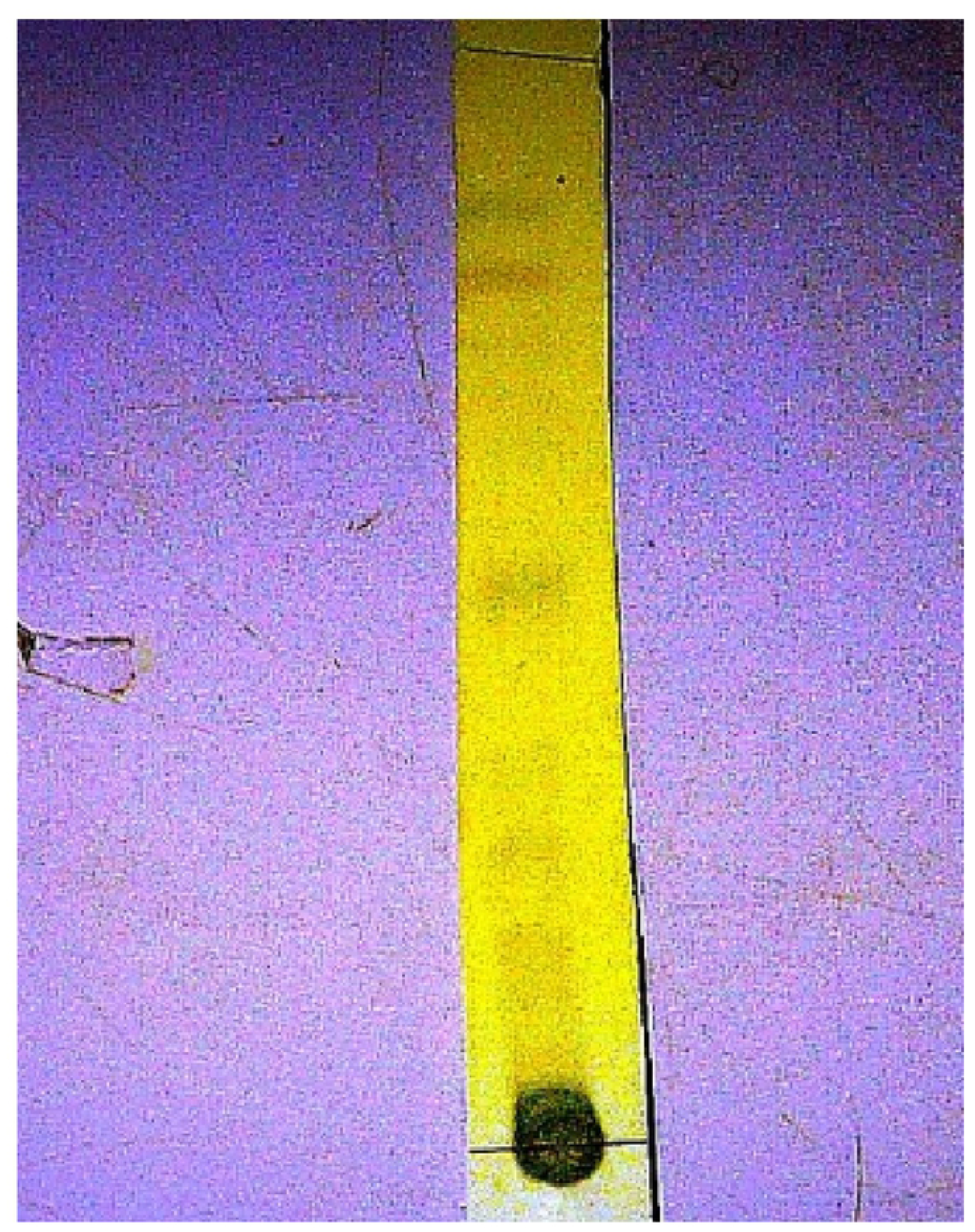
Thin layer chromatographic analysis of chloroform fraction (TBG-SK) sprayed with ferric chloride.

These spectra data therefore suggests a triterpenoid compound. This is supporting compound A to be rich in triterpenoids. To further support the presence of a triterpenoid, a thin layer chromatographic analysis of fraction A (Chlorofrm fraction) which produced compound A was carried out. The presence of triterpenoid compounds were indicated when sprayed with anisaldehyde in sulphuric acid reagent which produced a characteristic violet and purple coloured spots following activation of the plate “Fig 5”. Terpenoids-rich plants have been found useful in ethnomedicine, food, pharmaceutical and chemical industries [20].

Terpenoids are widely distributed in gymnosperms as well as in angiosperms and thus constitute a huge class of compounds germane to host plants for growth and developmental metabolism as well as for defense and to encourage pollination [21]. However, isoprenoid compounds of chemotaxonomic relevance biosynthesised by a limited group of flowering plants possess pharmacological properties important for drug development [22]. For instance, the tetracyclic triterpenoids have been reported in plant families including Rhamnaceae, Cucurbitaceae, Ganodermaceae and Apocynaceae while the pentacyclic triterpenoids are chemotaxonomically important in Rannunculaceae, Burseraceae, Capparidaceae, Celastraceae, Lamiaceae and Loranthaceae families of flowering plants [23]. As defense compounds terpenoids accumulate more in organs of the shoot system (such as the leaves and seeds) that are prone to pest attack or exposed to harsh environmental factors.

In the Loranthaceae angiospermic family, It was recently reported the isolation of seven new pentacyclic triterpenoids from the seeds of Tapinanthus banguensis which includes five oleanane-types designated bangwaoleanenes A–E, and two ursane-types, named bangwaursenes A and B. Other known related compounds isolated from this plant are: 3β-acetoxy-urs-12,13-ene-11-one, 3β-acetoxy-11α-hydroxyurs-12,13-ene, 11α,12α-oxidotaraxeryl acetate, β-amyrin acetate, (1R,5S,7S)-7-[2-(4-hydroxyphenyl)ethyl]-2,6-dioxabicyclo[3.3.1]nonan-3-one 1-desoxyribose, myo-inisitol, sorbitol, were isolated from the seeds of Tapinanthus bangwensis [12]. Generally, terpenoids have been reported to show a wide range of biopharmacological activities such as insecticidal, molluscicidal, fungicidal, spermicidal, anti-bacterial, analgesic, anti-viral, anti-inflammatory, anti-diabetic and cardiovascular [24], [25]. This may provide the basis for the varied ethnomedicinal uses of TBG in Nigerian South-western Traditional medicine where it is called “all-purpose plant” in folklore. In particular, sesquiterpenoids and triterpenoids are of scientific interest as potential sources of nature-derived angiogenic modulators [26]. The characterization of triterpenoids from the seeds of TGB [12] and our current isolation and detection of an anti-angiogenic triterpenoids in this plant is therefore consistent with previous literature reports. In summary, this study has provided scientific basis for the use of TBG-SK to modulate angiogenesis in Nigerian South-Western ethnomedicine.

## References

[1] Carmeliet P, Jain RK. Angiogenesis in cancer and other diseases. Nature. 2000;407: 249–257.

[2] Herrera A. In vivo evaluation of the potent angiosuppressive activity of some indigenous plants from Bataan, Philippines. Asia Life Sciences. 2010;19(1): 183–190.

[3] Folkman J. Angiogenesis in cancer, vascular, rheumatoid and other disease. Nature Medicine. 1995; 1: 27–31.

[4] Li WW, Li VW, Tsakayannis D. Emerging concepts and lessons from clinical trials of angiotherapy. The New Angiotherapy. 2001; 547–571.

[5] Ellis LM, Hicklin DJ. VEGF-targeted therapy: mechanisms of anti-tumour activity. Nature Reviews Cancer. 2008; 8: 579–91.

[6] Mahady GB. Global harmonization of herbal health claims. The Journal of nutrition. 2001; 131(3): 1120–1123.

[7] Xiaorui Z. World Health Organization-Traditional Medicine: Regulatory Situation of Herbal Medicines. A Worldwide Review. 1998; 1–5.

[8] Iwu MM. Hand book of African Medicinal Plants. 1993; 239.

[9] Arbonnier M. Trees, Shrubs and Lianas of West African Dry Zones. Cirad Margraf. 2004; 345–351.

[10] Kafaru E. Mistletoe an example of an all purpose herb herbal remedies. Guardian Newspaper. 1993; 11.

[11] Adesina SK, Illoh HC, Johnny II, Jacobs IE. African mistletoes (Loranthaceae); ethnopharmacology, chemistry and medicinal values: an update. African Journal of Traditional, Complementary and Alternative Medicines. 2013; 10(4): 161–170.

[12] Hermine M, Pierre M, Sandra LF, Sylvain KS, Hayato I, Hiroshi N, et al. Triterpenoids from seeds of Tapinanthus bangwensis. Phytochemistry letters. 2017; 19: 23–29.

[13] Nwafuru SK, Akunne TC, Ezenyi IC, Okoli CO. Anti-inflammatory Activity of Leaf Extract and Fractions of Tapinanthus bangwensis (Engl. & K. Krause) Danser Parasitic on Citrus angustifolia. European Journal of Medicinal Plants. 2017; 1–10.

[14] Oriola AO, Aladesanmi AJ, Arthur G. Anticancer activity of three African mistletoes. Nigerian Journal of Natural Products and Medicine. 2018; 22(1): 129–134.

[15] Trease GE, Evans WC. Pharmacognosy. Bailliere Tindall. 1985; 457–461.

[16] Muhammad Nihad AS, Rucha D, Vaijayanti PK, Ramesh RB, Savita PD. Establishment of an in ovo chick embryo yolk sac membrane (YSM) assay for pilot screening of potential angiogenic and anti-angiogenic agents. Cell Biology International. 2018; 1065–6995.

[17] Martin M, Bettina R, Bianca N, Weiwei X, Wen WR, Björn H, et al. Structure and hemodynamics of vascular networks in the chorioallantoic membrane of the chicken. J Physiol Heart Circ Physiol. 2016; 311(4): 913–926.

[18] Blake TA, Irina T, Jun OL, Charles MR. Itraconazole inhibits angiogenesis and tumor growth in non-small cell lung cancer. Cancer Research. 2011; 71(21): 6764–72.

[19] Stahl E. Thin Layer Chromatography. 2nd edition Springer and Academic Press. 1969; 123–366.

[20] Tholl D. Biosynthesis and biological functions of terpenoids in plants. In Biotechnology of isoprenoids. 2015; 63–106.

[21] Miettinen K, Inigo S, Kreft L, Pollier J, De Bo C, Botzki A, et al. The TriForC database: a comprehensive up-to-date resource of plant triterpene biosynthesis. Nucleic acids research. 2017; 46:586–594.

[22] Nguyen VT, Tung NT, Cuong TD, Hung TM, Kim JA, Woo MH, et al. Cytotoxic and anti-angiogenic effects of lanostane triterpenoids from Ganoderma lucidum. Phytochemistry letters. 2015; 12: 69–74.

[23] Bian X, Zhao Y, Guo X, Zhang L, Li P, Fu T, et al. Chiisanoside, a triterpenoid saponin, exhibits anti-tumor activity by promoting apoptosis and inhibiting angiogenesis. RSC Advances. 2017; 7(66): 41640–41650.

[24] Guan YY, Liu HJ, Luan X, Xu JR, Lu Q, Liu YR, et al. Raddeanin A, a triterpenoid saponin isolated from Anemone raddeana, suppresses the angiogenesis and growth of human colorectal tumor by inhibiting VEGFR2 signaling. Phytomedicine. 2015; 22(1):103–110.

[25] Guerra A, Duarte I, Duarte M. Role of Isoprenoid Compounds on Angiogenic Regulation: Opportunities and Challenges. Current medicinal chemistry. 2016; 23(9): 911–928.

[26] Park JH, Kim JK. Pristimerin, a naturally occurring triterpenoid, attenuates tumorigenesis in experimental colitis-associated colon cancer. Phytomedicine. 2018; 42:164–171.

